# Extrachromosomal circular DNA formation follows a 10.5-bp periodicity reflecting the minor groove orientation around nucleosomes

**DOI:** 10.64898/2026.10.19.760212

**Authors:** Aitor Zabala, Alex M. Ascensión, Frederik Otzen Bagger, Anders Krogh, David Otaegui, Iñigo Prada-Luengo

## Abstract

Extrachromosomal circular DNA is increasingly recognized as a central feature of somatic evolution in cancer, yet the structural determinants of eccDNA formation remain poorly understood. Here, we systematically analysed eccDNA length distributions and breakpoint organization across 25 datasets spanning multiple species, tissues and experimental approaches. Applying a short-time Fourier transform to eccDNA length distributions enabled us to resolve recurrent periodicities at approximately 180 bp and 10.5 bp, consistent with nucleosome length and the helical repeat of DNA, respectively. Mapping eccDNA breakpoints relative to nucleosome dyads revealed preferential breakpoint formation in linker DNA and, within nucleosome-bound DNA, at positions where the minor groove faces towards the histone surface. Our results show that eccDNA formation is shaped not only by nucleosome positioning but also by the rotational orientation of the DNA double helix around the nucleosome. Together, these findings identify chromatin geometry as a determinant of eccDNA biogenesis and provide a structural basis for the length periodicities that characterize eccDNA formation.

## Introduction

Extrachromosomal circular DNA (eccDNA) is an abundant and heterogeneous component of the genome, consisting of chromosomally derived DNA circles that exist independently of chromosomes (Cohen and Mechali, 2001; Cohen et al., 2008; Møller et al., 2015, 2018, 2020). EccDNA are widespread across a broad range of eukaryotic species and cellular contexts, including normal cells, ageing, and cancer (Cohen et al., 2003; Møller et al., 2015; Turner et al., 2017; Møller et al., 2018; Kim et al., 2020; Prada-Luengo et al., 2020; Liang et al., 2025). Rather than representing incidental by-products of genomic instability, eccDNAs are now recognized as recurrent components of the nuclear DNA landscape with potential functional consequences, such as oncogene amplification and remodelling (Nathanson et al., 2014; Morton et al., 2019; Helmsauer et al., 2020). EccDNA can arise through DNA double-strand break (DSB) associated excision and circularization of chromosomal sequences (Cohen and Lavi, 2009; Møller et al., 2018). Depending on the genomic locus and cellular context, circularization can involve canonical non-homologous end joining or alternative end joining, including microhomology-mediated end joining (Cohen et al., 2006; Paulsen et al., 2021; Rose et al., 2025; Chung et al., 2026). EccDNA molecules often exhibit length distributions corresponding to the length of the nucleosome, with prominent peaks around 180 bp and higher-order multiples (Shibata et al., 2012; Dillon et al., 2015; Møller et al., 2018; Paulsen et al., 2018; Kumar et al., 2020; Sin et al., 2020, 2021, 2022; Lv et al., 2022; Hu et al., 2023; Ling et al., 2024; Wang et al., 2024). In addition to these nucleosome-scale features, circulating cell-free eccDNA and nuclease-accessibility-based datasets have reported finer-scale periodic signatures on the order of 10 bp, associated with nuclease cleavage during apoptosis (Sin et al., 2020, 2022).

However, how nucleosome organization and DNA helical geometry contribute to the periodic patterns observed in eccDNA length distributions remains unresolved. The double helix exhibits an helical periodicity of approximately 10.5 bp per turn (Klug and Lutter, 1981). As DNA wraps around the histone octamer, its rotational orientation causes the minor groove to face alternately towards and away from the histone surface at approximately half-helical turn intervals (Klug and Lutter, 1981). Genome-wide analyses have shown that mutation patterns within nucleosome-occupied DNA can display an approximately 10-bp periodicity associated with this rotational organization (Pich et al., 2018), while similar periodic patterns have been observed in UV-induced DNA damage and the accessibility of break-induced replication repair enzymes (Gale et al., 1987; Hinz et al., 2010; Mao et al., 2016). Whether DNA rotational orientation influences eccDNA breakpoint formation, still, remains unknown. We investigated if eccDNA length distributions contain a signature of the 10.5-bp helical repeat and whether nucleosome positioning and the rotational orientation of nucleosomal DNA contribute to eccDNA formation.

In our study, we analyse eccDNA length distributions across 25 datasets spanning multiple species, biological samples, and experimental platforms. By applying spectral decomposition to eccDNA length profiles, we reveal a recurrent periodicity of approximately 10.5 bp across diverse species and experimental protocols, consistent with the helical repeat of DNA. We further investigate the structural basis of this periodicity by examining eccDNA breakpoint positions relative to nucleosome organization. We show that eccDNA breakpoints are preferentially enriched in linker DNA and, within nucleosome-bound DNA, exhibit a rotational preference for positions where the minor groove faces toward the histone surface. Together, these findings reveal that eccDNA formation is shaped by both nucleosome positioning and the rotational organization of nucleosomal DNA.

## Results

EccDNA molecules often exhibit non-random length distributions, with prominent peaks around 180 bp and higher-order multiples (Shibata et al., 2012; Dillon et al., 2015; Møller et al., 2018; Sin et al., 2020; Yang et al., 2021; Sin et al., 2022; Lv et al., 2022; Hu et al., 2023). These peaks support a contribution of nucleosome-associated organization in eccDNA formation. In addition to these long-period features, some studies have observed shorter periodic signatures of 10 bp (Sin et al., 2020, 2022), although their biological significance and relationship to eccDNA biogenesis remain largely unresolved. We reasoned that such short-period signals could reflect the underlying helical geometry of double-stranded DNA (**Figure 1A**). To test this hypothesis, we systematically explored eccDNA lengths to determine whether short-period structure could be detected beyond the broad nucleosome-associated peaks (**Figure 1B**), and whether such signals were conserved across biological contexts and experimental approaches. We analysed 25 datasets spanning multiple species, including *Homo sapiens*, *Mus musculus*, *Gallus gallus*, and *Arabidopsis thaliana*, and a range of tissues such as blood, plasma, muscle, brain, and immune and endocrine organs. To explore whether the signals were consistent across experimental approaches, we included datasets generated using different eccDNA enrichment strategies: microDNA-seq (Shibata et al., 2012), Circle-Seq (Møller et al., 2016), ATAC-seq-derived eccDNA (Sin et al., 2020), and eccDNA-seq approaches (Zhang et al., 2021) (**Figure 1C**; **Table 1**). Because not all datasets were originally generated or processed using a uniform eccDNA detection approach, we reprocessed all sequencing data using a uniform bioinformatic workflow (Schreyer et al., 2024), which implements Circle-Map for eccDNA calling, to obtain nucleotide-resolution eccDNA coordinates. Finally, to ensure high-confidence eccDNA calls, only molecules supported by at least two split reads were included in downstream analyses. Altogether, this yielded 9,640,449 high-confidence eccDNAs across the 25 datasets with a mean of 385,618 eccDNAs per sample (standard deviation = 1,657,292), which were used for downstream analysis.

**Figure 1.**
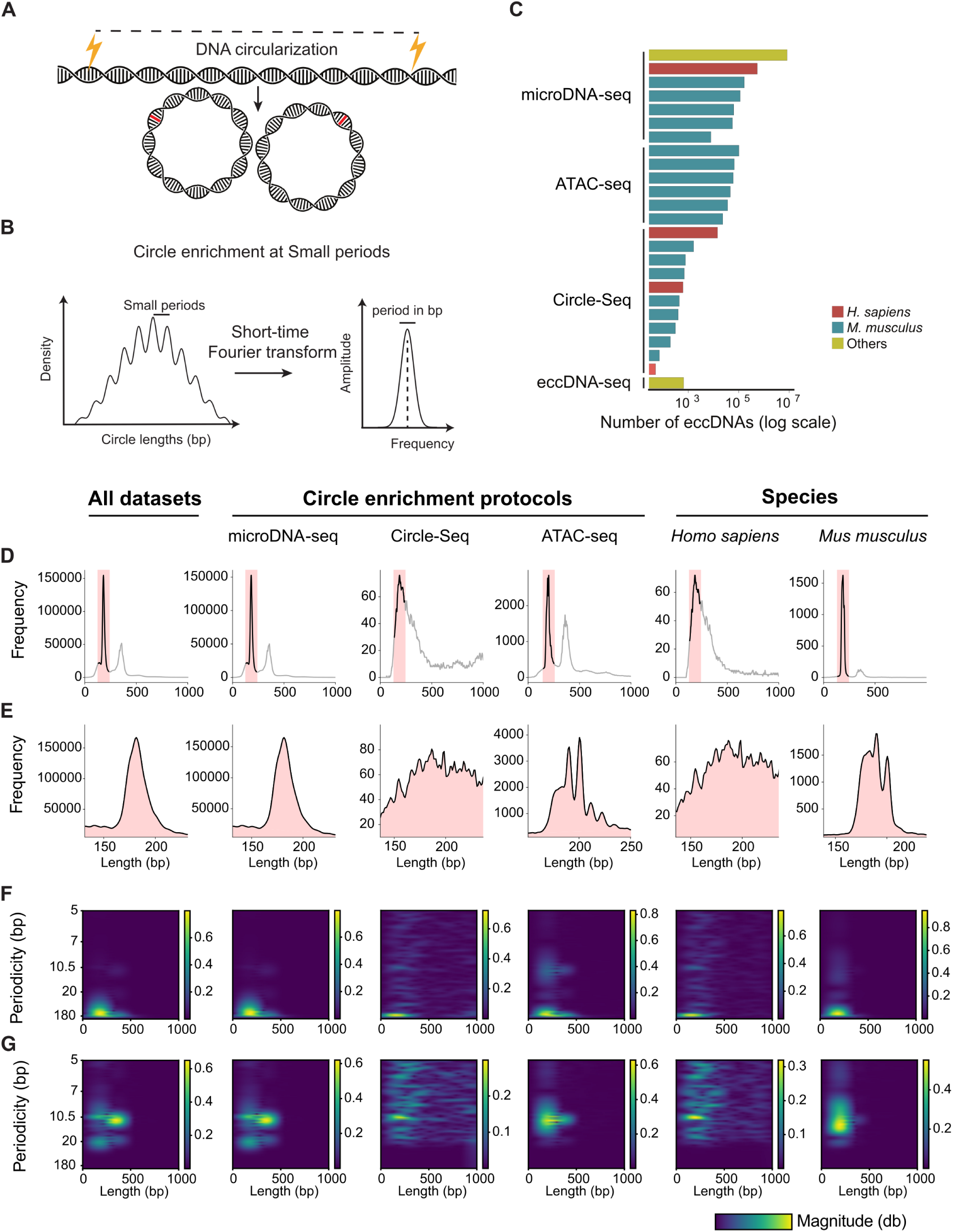
eccDNA length distributions reveal multi-scale periodicity. (**A**) Conceptual model of eccDNA formation through double-strand breaks (DSBs) followed by circularization. The schematic highlights the expected 10.5-bp periodicity (high-frequency signal), corresponding to the helical geometry of double-stranded DNA, which may influence preferred sites of DNA cleavage and loop formation. (**B**) Computational framework for detecting multi-scale periodicity. EccDNA length distributions are analysed using STFT to resolve both 180 bp and 10.5 bp periodicities associated with nucleosome organization and DNA helical geometry, respectively. (**C**) Distribution of dataset sizes. Bar plot showing the total number of eccDNAs per dataset (log scale), highlighting variability in sample size and its impact on the detection of short-length periodicity. (**D**) Global eccDNA length distributions up to 1000 bp, showing dominant and secondary peaks across datasets. (**E**) Zoomed view of the dominant peak region, revealing subtle short-length oscillatory patterns. (**F**) STFT analysis showing global periodicity profiles obtained using a Butterworth high-pass filter (cutoff = 0.005), capturing the 180-bp periodicity (low-frequency signal) associated with nucleosome organization. (**G**) STFT analysis focused on local periodicity profiles obtained using a higher cutoff frequency (0.05), capturing the 10.5-bp periodicity (high-frequency signal) associated with DNA helical geometry.

**Table 1.**
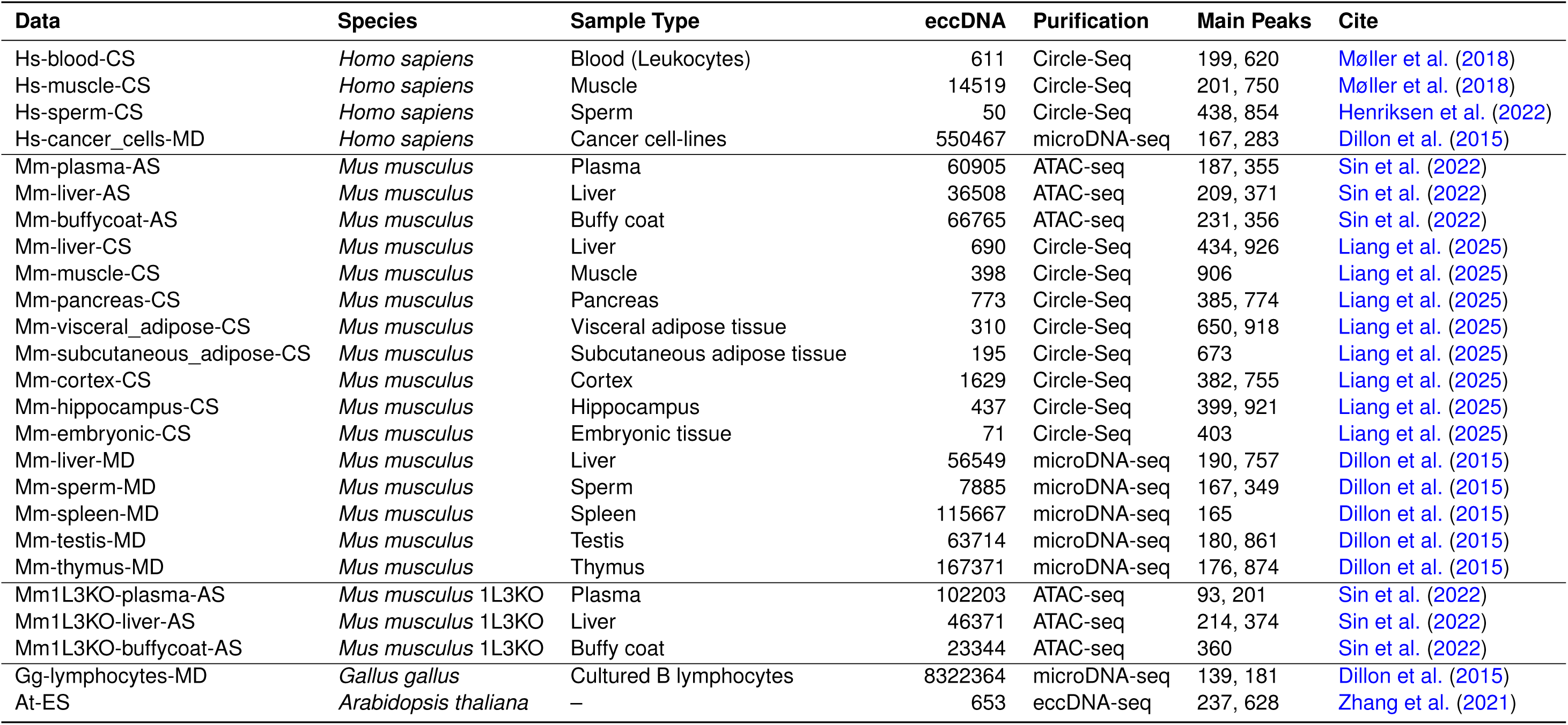
Summary of datasets by organism, sample type, purification method, eccDNA count, and main eccDNA length peaks.

We focused on short eccDNA molecules (eccDNA smaller than 1000 bp) to characterize their length distributions, as these represented the vast majority of detected eccDNAs. As expected, length profiles showed a prominent enrichment around 180 bp, often accompanied by a secondary peak near 360 bp (**Figure 1D**). This 180-bp peak was observed in 16 of 25 datasets, indicating that nucleosome-scale enrichment is a recurrent feature of short eccDNA length distributions. We further confirmed that these peaks were present across datasets generated using different experimental protocols, including ATAC-seq-derived eccDNA, microDNA-seq, and Circle-Seq and different species (**Figure 1D** & **Supplementary Figure S1**). Interestingly, computational peak detection revealed that the detected peaks were not uniformly centred at 180 bp: 15 of the 25 datasets showed dominant peaks between approximately 165 and 245 bp (**Table 1**), a range that overlaps the nucleosome repeat-length scale and can be generated by a relatively invariant 147-bp nucleosomal core together with variable linker DNA (Kornberg, 1974; Widom, 1992). Notably, the *Hs-cancer_cells-MS* dataset exhibited a peak at 167 bp, a short nucleosome repeat length previously used in reconstituted chromatin arrays (Routh et al., 2008). Although this correspondence does not establish that the eccDNA molecules directly recapitulate nucleosome repeat length, it is consistent with a contribution of nucleosome organization to eccDNA formation. Furthermore, the enrichment became more apparent when datasets were grouped according to the method used for eccDNA purification, with peaks centred at approximately 181 bp for microDNA, 207 bp for Circle-Seq, and 193 bp for ATAC-seq-derived eccDNA. Together, these observations support a recurrent nucleosome-scale enrichment in eccDNA length distributions, while indicating that its precise position varies across datasets and experimental approaches.

We next explored whether finer-scale periodic patterns were present around the dominant peak. To account for variation in the 180-bp peak position across datasets, we centered each length distribution on its mode and examined a 100-bp window around this point. Within this window, we observed weak but consistent oscillations at approximately 10-bp intervals across most groups (**Figure 1E**). This pattern was present in both Circle-Seq and ATAC-seq-derived datasets and across multiple tissues and species. However, no clear 10-bp periodic signal was apparent in the *Arabidopsis thaliana* sample At-ES, the *Gallus gallus* dataset, or samples generated using microDNA-seq (**Figure 1E** & **Supplementary Figure S1**). Together, these observations suggested the presence of an approximately 10-bp periodic signal, although raw length distributions alone made it difficult to resolve it confidently and assess its extent across datasets and species.

To characterize these periodic signals more systematically, we performed a Fourier analysis, which provides a sensitive and objective means of detecting weak oscillatory components that are not readily apparent in raw length distributions. The classic Fourier transform, however, collapses the entire length spectrum into a single global frequency representation and cannot determine whether a periodic signal is restricted to a particular eccDNA size range. We therefore applied a short-time Fourier transform (STFT), which estimates frequency content locally across the eccDNA length spectrum, allowing periodic components to be resolved while retaining information about the length ranges in which they appear (**Figure 1B** and Materials and Methods). The STFT revealed a pronounced periodic component around 180 bp across datasets, experimental protocols, and species (**Figure 1F**), with *Arabidopsis thaliana* as the only exception (**Supplementary Figure S2**). The 180-bp signal was visually clear in all samples containing more than 7,885 eccDNAs, representing 14 of 25 datasets (**Supplementary Figure S2**).

We next focused specifically on the putative shorter periodicity. Even before frequency filtering, a component around 10.5 bp was visible in several STFT profiles, particularly in ATAC-seq-derived datasets and human datasets (**Figure 1F**), and many individual datasets (**Supplementary Figure S2**). To isolate high-frequency components corresponding to short-period oscillations, we applied a Butterworth high-pass filter retaining periodicities up to 20 bp. This revealed a prominent signal centered at a periodicity of approximately 10.5 bp across datasets and experimental protocols (**Figure 1G** and **Supplementary Figure S2**). Notably, the signal was also detected in *Gallus gallus* and in the microDNA-seq and “All datasets” groups, although it was not readily apparent in their raw length histograms, highlighting the sensitivity of the Fourier-based analysis for resolving weak periodic patterns. As with the 180-bp component, *Arabidopsis thaliana* was the only species in which the 10.5-bp periodicity was not detected.

Despite the sensitivity of STFT to detect complex signals that barely went unnoticed with the naked eye, the characteristic periodicity of 10.5 bp was not clearly detected in 11 of 25 datasets. We noted that all datasets lacking a detectable peak contained relatively few eccDNAs and that dataset size clearly separated samples according to whether the peak was detected (**Supplementary Figure S2**, Mm-sperm-MD sample), suggesting that the signal may have gone undetected because of limited statistical power rather than being biologically absent. To assess the effect of dataset size on signal detection, we compared the maximum STFT magnitude at the 10.5-bp periodicity with the number of eccDNAs in each dataset (**Supplementary Figure S3**). The magnitude increased with dataset size: it remained low in datasets containing fewer than *∼* 10^3^ eccDNAs and rose markedly in datasets above *∼* 10^4^ eccDNAs, suggesting that the 10.5-bp periodicity is not reliably detectable in datasets containing fewer than approximately 10,000 circles. Together, these findings support the presence of an approximately 10.5-bp periodic signal in eccDNA length distributions across diverse biological and experimental contexts. The reproducibility of this signal suggests that it is unlikely to be explained solely by a specific eccDNA enrichment or detection protocol, although its biological origin remains unclear.

To investigate the structural basis of this periodicity, we considered two candidate hypotheses: the intrinsic helical rotation of the DNA double helix, and the alternating inward- and outward-facing orientation of the minor groove as DNA wraps around the nucleosome, as previously reported by others for single nucleotide variation (Pich et al., 2018). We first examined whether eccDNA molecules showed a preferred phase with respect to the 10.5-bp helical repeat. For each circle, we calculated its phase within a 10.5-bp cycle based on its length (Material and Methods). If eccDNA lengths preferentially occurred at exact multiples of the helical repeat, we would expect an enrichment around phases that favour circle closure, in other words, around 0*^◦^* and 360*^◦^*. The phase distribution showed no pronounced peak at 0^◦^ or 360*^◦^*, and more generally no strong preference for a particular phase (**Supplementary Figure S4A**). Thus, the observed 10-bp periodicity does not appear to reflect a preference for circle lengths corresponding to exact multiples of the DNA helical repeat.

We next investigated whether the 10-bp periodicity could be related to the positioning of eccDNA breakpoints relative to nucleosomes. To this end, we obtained MNase-seq data (Gaffney et al., 2012) and mapped nucleosome positions across the genome. To define a high-confidence set of nucleosomes, we quantified their translational stability based on the sharpness of the MNase-seq signal and restricted subsequent analyses to translationally stable nucleosomes (positioning score *≥* 0.5; Material and Methods). We further focused on intergenic eccDNA to minimize variability associated with nucleosome organization within transcriptionally active regions (Lee et al., 2004; Pich et al., 2018). For each eccDNA breakpoint, we calculated its position relative to the nearest nucleosome dyad (the central base pair of the nucleosomal DNA, located on the nucleosome’s axis of pseudo-two-fold symmetry), using consensus dyad positions inferred from MNase-seq data (Material and Methods), and determined observed-to-expected breakpoint enrichment by comparison with 1000 randomized eccDNA coordinates. Breakpoints showed a clear enrichment at the linker DNA, extending from the dyad towards its outer boundaries (**Figure 2A**). Enrichment was particularly pronounced at *±*83.5 bp from the dyad, defining a symmetric interval of approximately 167 bp around the nucleosome center, corresponding to the sum of the nucleosome and linker DNA length. Consistent with this boundary-associated pattern, eccDNA breakpoints were approximately 1.6-fold enriched in linker regions relative to the randomized controls. Together, these results indicate that eccDNA breakpoints are non-randomly positioned with respect to nucleosome architecture, with a strong preference for the nucleosome boundary and adjacent linker DNA.

**Figure 2.**
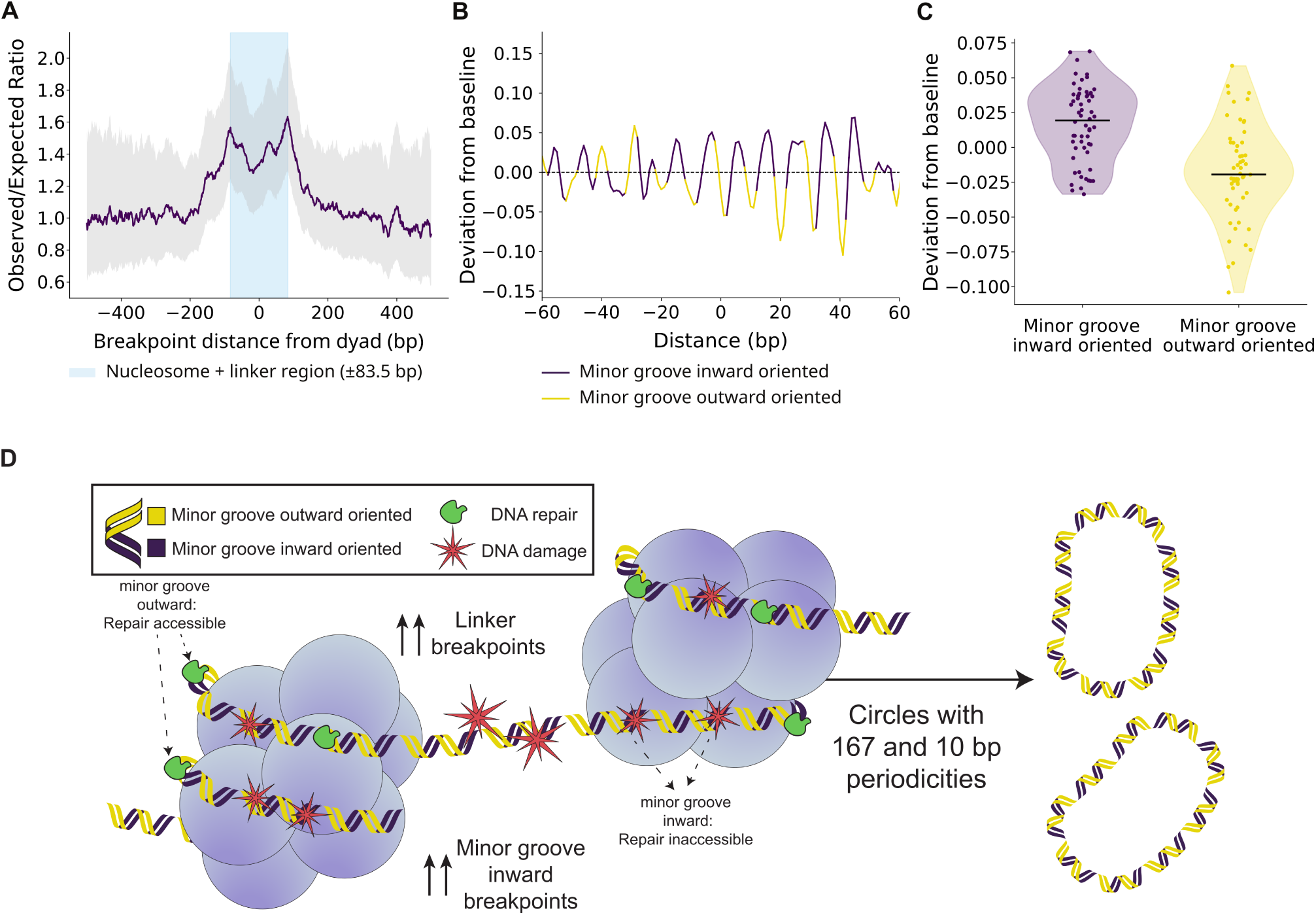
Consensus nucleosome dyads define the spatial pattern and orientation of eccDNA circle breakpoints. (**A**) Observed/expected ratio of eccDNA breakpoint positions relative to the nearest nucleosome dyad. The shaded region indicates the nucleosome and linker region (*±*83.5 bp), revealing increased breakpoint enrichment near the nucleosome boundaries and depletion around the dyad. (**B**) Deviation from the baseline profile, highlighting periodic differences associated with minor groove orientation. Purple and yellow indicate minor groove inward- and outward-oriented positions, respectively. (**C**) Distribution of deviations from baseline at minor groove positions, separated by groove orientation. Minor groove inward-oriented positions (*n* = 60) showed significantly higher deviations than outward-oriented positions (*n* = 55) (Welch’s *t*-test, *t* = 5.96, *p* = 3.73 *×* 10*^−^*^8^; Cohen’s *d* = 1.12). (**D**) Proposed model for how nucleosome architecture and DNA rotational geometry jointly shape eccDNA formation. DNA is coloured by the rotational orientation of the minor groove (purple, facing inward towards the histone octamer; yellow, facing outward). DNA damage (red) arising in linker DNA is readily accessible to repair factors (green), whereas within nucleosome-bound DNA accessibility depends on rotational phase: outward-facing lesions remain accessible, while inward-facing lesions are occluded by the histone surface and are processed less efficiently. Breakpoints therefore accumulate preferentially in linker DNA and, within the nucleosome, at inward-facing minor-groove positions, giving rise to circles whose length distributions carry the nucleosome-plus-linker (*∼*167 bp) and DNA helical-repeat (*∼*10.5 bp) periodicities.

Having identified a broader nucleosome-associated enrichment, we asked whether breakpoint positioning also reflected the rotational geometry of nucleosomal DNA. As DNA wraps around the histone octamer, its rotational orientation changes periodically along the superhelical locations (SHLs), such that the minor groove alternates between facing toward and away from the histone surface. Positions at approximately *±*5, *±*15, *±*25, and subsequently every *∼* 10 bp up to *±*65 bp from the dyad correspond to regions where the minor groove is oriented inward, facing towards the histones. Previous studies have reported that the rotational orientation of nucleosomal DNA is associated with differences in the frequency of somatic single-nucleotide mutations (Pich et al., 2018). We therefore examined the breakpoint enrichment profile at higher resolution around the dyad (**Supplementary Figure S4B**). This showed a clear oscillatory pattern, with local enrichment peaks recurring at approximately 10-bp intervals. Notably, these peaks frequently coincided with positions where the minor groove faces toward the histone surface, suggesting that eccDNA breakpoint formation may be sensitive to the rotational orientation of nucleosomal DNA.

This fine-scale oscillation, however, was superimposed on the broader increase in breakpoint enrichment toward the nucleosome boundaries, making the two effects difficult to distinguish directly. To isolate the rotational component, we removed this broad nucleosome-associated trend using Savitzky-Golay smoothing (Savitzky and Golay, 1964). We estimated the global enrichment trend using a 21-bp window and the local minor-groove-scale variation using a shorter 5-bp window, allowing us to assess the relative local increase in eccDNA breakpoints above the underlying nucleosome-scale baseline. After this correction, inward-facing minor-groove positions tended to lie above the local baseline (**Figure 2B**). In fact, 10 of the 12 local enrichment peaks coincided with positions at which the minor groove is oriented inward, toward the histones. At nucleotide resolution, this rotational preference became more apparent, with inward-facing positions showing a consistent relative increase in breakpoint enrichment compared with outward-facing minor-groove positions (**Figure 2C**). We quantified this difference by comparing residual enrichment values extracted around the two classes of rotational positions (**Figure 2C**). Minor-groove-inward oriented positions showed higher residual values (median = 0.0194, *n* = 60) than minor-groove-outward oriented positions (median = *−*0.0195, *n* = 55), showing a significant difference (*t* = 5.96, *P* = 3.73 *×* 10*^−^*^8^; Cohen’s *d* = 1.12). Together, these results show that eccDNA breakpoints have a rotational preference for positions where the minor groove is oriented inward.

## Discussion

Understanding the mechanisms that govern eccDNA formation is key to clarifying its origins and potential contributions to genome plasticity and adaptation. Here, we show that eccDNA breakpoints are preferentially located in linker DNA resulting in eccDNA with preferential lengths around the 180bp and 360bp peaks, and within nucleosome-bound DNA, at positions where the minor groove faces towards the histone surface. This preference for inward-facing minor-groove positions supports nucleosomal DNA geometry as a structural basis for the approximately 10.5-bp periodicity observed in short eccDNA length distributions. Together, these findings support a model in which DNA rotational orientation complements nucleosome positioning in shaping short eccDNA formation (**Figure 2D**).

We detected a periodicity of approximately 10.5 bp in eccDNA length distributions across datasets containing sufficient circles to resolve this fine-scale signal. Sin et al. (2020) linked a 10-bp periodicity of cell-free eccDNA to nucleosomal organization and demonstrated that extracellular nuclease activity shapes plasma eccDNA size distributions. Here, we show that this periodicity recurs across a much broader range of species, tissues, biological conditions and experimental protocols, extending its relevance beyond circulating cell-free DNA (Sin et al., 2020, 2022). Together with the preferential localization of breakpoints at inward-facing minor-groove positions, this broader occurrence supports nucleosomal DNA geometry as a structural influence on eccDNA biogenesis. The inward-facing preference is initially counterintuitive, because these positions are less accessible to solvent-exposed DNA-binding factors and some DNA-damaging agents (Gale et al., 1987; Mao et al., 2016). However, the orientation of the DNA groove governs not only where damage occurs, but also how lesions are recognized and repaired. Inward-facing lesions are generally less accessible to DNA repair factors because they are occluded by the histone surface; repair proteins may therefore need to reposition the nucleosomal DNA to access lesions facing the histone core (Matsumoto et al., 2019). More broadly, the rotational and translational positions of DNA lesions influence their accessibility to base-excision repair pathway enzymes (Hinz et al., 2010; Rodriguez et al., 2015). Consistent with this mechanism, lesions oriented toward the histone surface can be processed less efficiently than those oriented away from it, particularly near the nucleosomal dyad, where outward-facing uracils show substantially higher excision rates in nucleosome core particles (Cole et al., 2010; Rodriguez et al., 2015). Similar inward-facing mutation patterns have been reported in association with *POLE* deficiency and oxidized guanine incorporation (Pich et al., 2018), as well as in esophageal adenocarcinoma (Morledge-Hampton and Wyrick, 2021). Altogether, these observations suggest that the rotational phase of eccDNA breakpoints may reflect the underlying mechanisms of DNA damage recognition and repair, rather than DNA accessibility alone.

We also tested whether the intrinsic helical geometry of DNA favours circularization at particular lengths by examining eccDNA length phases relative to the 10.5-bp helical repeat. We found no pronounced phase preference, arguing against a simple rotational-alignment model for circle length. However, context-dependent effects of helical geometry on closure, as observed in DNA cyclization experiments (Shore et al., 1981; Shore and Baldwin, 1983), remain possible. A separate consideration is amplification bias: Joffroy et al. (2018) demonstrated a similar helical-repeat-associated periodicity in *ϕ*-29 mediated rolling-circle amplification of very short circular templates. However, our detection of the signal in *ϕ*-29 free ATAC-seq-derived datasets argues against *ϕ*-29 amplification as its sole source. Its persistence at eccDNA lengths approaching 500 bp further weakens an explanation based exclusively on this short-template effect. Collectively, these findings connect recurrent length periodicity to nucleosome-relative breakpoint organization, supporting a structural contribution from DNA rotational positioning while leaving open additional effects of DNA geometry on circularization.

Using nucleosome maps from human lymphoblastoid cell lines, we identified a preferential enrichment of eccDNA breakpoints in linker DNA, with pronounced peaks at 83.5 bp on either side of the nucleosome dyad. This pattern is consistent with previous evidence of elevated double-strand breaks around translationally stable nucleosomes (Li and Luscombe, 2020). Nucleosome-associated origins of eccDNA have long been proposed (Hu et al., 2023; Sin et al., 2020; Shibata et al., 2012). Our findings extend these observations by identifying preferred breakpoint positions relative to nucleosome dyads, providing positional support for a nucleosome-associated explanation of the recurrent approximately 180 bp eccDNA length peak (Shibata et al., 2012; Dillon et al., 2015; Møller et al., 2018; Paulsen et al., 2018; Kumar et al., 2020; Sin et al., 2020, 2021, 2022; Lv et al., 2022; Hu et al., 2023; Ling et al., 2024; Wang et al., 2024). However, although 15 of the 25 datasets reported a dominant eccDNA length peak between 165 and 245 bp, the position of this peak was not uniform across datasets. This range overlaps the nucleosome repeat-length scale, which comprises the relatively invariant 147 bp of DNA wrapped around the histone octamer together with a variable linker-DNA segment (Kornberg, 1974; Widom, 1992). In particular, the 167-bp peak observed in the *Hs-cancer_cells-MS* dataset is compatible with a short nucleosome repeat length previously used in reconstituted chromatin arrays (Routh et al., 2008). Thus, the enrichment of breakpoints at approximately ±83.5 bp from the nucleosome dyad provides a positional explanation for the shorter eccDNA peaks in this interval, whereas longer peaks may reflect additional linker DNA and differences in local nucleosome spacing.

Preferential accessibility near nucleosome boundaries could favour cleavage and fragment release at these locations, generating nucleosome-scale precursors for circularization. A major limitation, however, is that the nucleosome and eccDNA maps were not obtained from the same biospecimens. To mitigate potential differences in nucleosome organization, we selected translationally stable nucleosomes and restricted the analysis to intergenic eccDNA, aiming to minimize variability associated with transcriptionally active regions. We envision that future studies with eccDNA and nucleosome maps from the same biospecimen will provide a much cleaner picture of the signal. Future studies could also experimentally reposition nucleosomes on the same DNA sequence and test whether eccDNA breakpoint enrichment shifts with nucleosome boundaries. A corresponding shift would support a causal contribution of nucleosome positioning to eccDNA formation, rather than a preference determined solely by the underlying DNA sequence. Our results move beyond interpreting length peaks alone by establishing where eccDNA breakpoints preferentially occur relative to nucleosomes.

Our results support a model of eccDNA formation in which nucleosome positioning and DNA rotational orientation jointly shape the observed length distributions and breakpoint patterns (**Figure 2D**). EccDNA breakpoints are preferentially enriched in linker regions, consistent with greater accessibility relative to nucleosome-bound DNA. Within nucleosomes, however, breakpoint positioning follows the rotational orientation of the DNA, with increased enrichment at inward-facing minor-groove positions. This pattern suggests that rotational positioning may influence the processing and repair of DNA damage, rather than simply determining where damage occurs. Together, these findings place nucleosomal architecture and DNA rotational geometry as complementary structural constraints on eccDNA biogenesis, with potential implications for understanding genome-editing outcomes and drug–DNA interactions in chromatin.

## Methods

### Biological datasets

We analysed a total of 25 biological datasets derived from four major multicellular eukaryotic reference species: *Homo sapiens*, *Mus musculus*, *Gallus gallus*, and *Arabidopsis thaliana*. These datasets encompass a diverse range of biological samples, including blood, plasma, buffy coat, sperm, muscle, liver, brain regions such as cortex and hippocampus, immune and endocrine-associated tissues including thymus, spleen, testis, pancreas, and adipose tissue, as well as cancer cell lines.

Across these studies, eccDNA was isolated or enriched using a variety of approaches, including Circle-Seq (Møller et al., 2016), ATAC-seq-based enrichment of cell-free extrachromosomal DNA (Sin et al., 2020), selective enzymatic digestion of linear DNA followed by targeted amplification to isolate short extrachromosomal DNA, defined as microDNA-seq in this study (Shibata et al., 2012), and eccDNA-seq (Zhang et al., 2021), a term used by the original authors to collectively describe Mobilome-seq (Lanciano et al., 2021), Circle-Seq, and CIDER-seq (Mehta et al., 2020).

For *Homo sapiens*, we included four datasets: blood, muscle, and sperm samples purified using Circle-Seq, and cancer cell lines generated using microDNA-seq described above. *Mus musculus* was represented by sixteen datasets: plasma, liver, and buffy coat samples generated using ATAC-seq; liver, muscle, pancreas, visceral adipose tissue, subcutaneous adipose tissue, cortex, hippocampus,and embryonic tissue using Circle-Seq; and liver, sperm, spleen, testis, and thymus samples generated using microDNA-seq. We also collected ATAC-seq datasets from plasma, liver, and buffy coat of knockout mouse models deficient in deoxyribonuclease 1-like 3 (DNASE1L3). For *Arabidopsis thaliana*, we used a dataset generated using eccDNA-seq. An overview of the sample type, species, and purification method is provided in **Table 1**.

### eccDNA detection

To ensure homogeneous and consistent computational processing across datasets, we downloaded and processed all raw sequencing data using Circle-Map v.1.1.4 (Prada-Luengo et al., 2019) within the nf-core/circdna pipeline v1.1.0 (Schreyer et al., 2024). Briefly, the pipeline maps the reads with BWA-MEM (v0.7.17-r1188) (Li and Durbin, 2009; Li, 2013):, sorts & indexes SAM files with SAMtools v1.16.1 (Li et al., 2009; Danecek et al., 2021): to then proceed with Circle-Map. We selected Circle-Map because its nucleotide-level resolution of eccDNA breakpoint coordinates enables accurate characterization of the eccDNA frequency spectrum. Circle-Seq datasets were excluded from this reprocessing step because the original authors processed them using Circle-Map v1.1.4. The reference genomes provided to the nf-core/circdna pipeline were GRCh38 for *Homo sapiens*, GRCm38 for *Mus musculus*, Galgal4 for *Gallus gallus*, and TAIR10 for *Arabidopsis thaliana*. We downloaded all the Circle-Seq data originating *Homo Sapiens* and *Mus musculus* from the supplementary materials of the original publications (Møller et al., 2018; Henriksen et al., 2022; Dillon et al., 2015; Liang et al., 2025).

### Short-time Fourier transform

The biogenesis of eccDNAs may be shaped by underlying structural features of chromatin and DNA, such as nucleosome organization and DNA helical periodicity. However, these patterns are often obscured in the raw length-density signal and are difficult to detect from histograms alone. To uncover and characterize these hidden periodic components, we analyzed the density signal in the frequency domain, with the aim of identifying the frequencies that contribute to the observed length distribution.

Let *f* (*x*) denote the eccDNA length-density signal, where *x* is eccDNA length in base pairs (treated as a real number). The Fourier transform decomposes this signal into sinusoidal components of different frequencies,

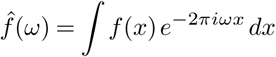

where *ω* denotes frequency in cycles per base pair. A peak at frequency *ω* corresponds to a periodic component with period 1*/ω* base pairs. This allows us to identify which periodicities contribute to the observed length distribution, for example periodic components compatible with nucleosome spacing or DNA helical periodicity. To describe how localized the signal is in length space, the mean and variance of the length are defined as:

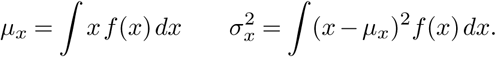

The mean frequency and frequency variance are defined from the Fourier-transformed signal as

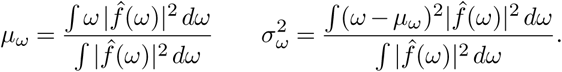

With these quantities defined, the Fourier uncertainty principle states that the signal cannot be arbitrarily localized in both length and frequency space; equivalently, the product of the two standard deviations is bounded below by

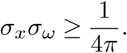

This implies that a signal cannot be sharply localized in both eccDNA length and frequency at the same time. In practice, this creates a trade-off: strong localization along the length axis reduces frequency resolution, whereas high frequency resolution requires averaging over a broader range of lengths. For example, determining whether a 10.5-bp DNA helical periodicity is present at a specific eccDNA length requires strong localization along the length axis, but this comes at the cost of reduced precision in estimating its frequency. Conversely, estimating the 10.5-bp periodicity with high frequency precision requires averaging over a broader range of eccDNA lengths, making it harder to determine exactly where the signal is active. This is particularly relevant because we expect the DNA helical signal to be non-stationary, meaning that its periodic structure may change across length and decay with increasing eccDNA length.

To obtain this positional information, we used the short-time Fourier transform (STFT) (Oppenheim and Schafer, 2009), which applies the Fourier transform locally within an sliding window. For any non-negative window function *w*(*x−τ* ) with finite support, centered at eccDNA length *τ* , the STFT is defined as

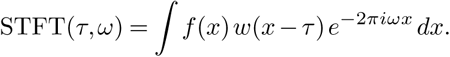

Here, *τ* determines the position of the window along the length axis, while *ω* denotes frequency. The resulting function therefore describes the frequency content of the density signal locally as a function of eccDNA length, allowing the detection of localized periodic components that are not identifiable under global spectral decomposition.

The STFT was computed using the scipy.signal.stft with a Hann window (nperseg = 600, noverlap = 599, fs = 1.0), focusing on frequencies between 0 and 0.20 Hz. The choice of window length (600 bp) reflects a trade-off between frequency resolution and local sensitivity along the length axis, while the high overlap (599 bp) ensures maximal spatial resolution, effectively yielding a sliding-window analysis with single–base-pair steps. The Hann window minimizes spectral leakage and reduces boundary artifacts associated with finite-length segments (Oppenheim and Schafer, 2009), ensuring smooth transitions between adjacent windows and improving the robustness of local spectral estimates.

The STFT output is a complex spectrogram representing how the spectral content of eccDNA length distributions varies over length and frequency. The *x*-axis corresponds to position along the length distribution, and the *y*-axis corresponds to frequency (inverse base pairs). Amplitudes, *|STFT* (*τ, ω*)|, were color-coded to highlight regions of significant local periodicity.

### Preprocessing

For preprocessing, we converted eccDNA length distributions into length-density histograms, followed by filtering to retain only eccDNAs smaller than 1000 bp. We applied this filtering step to focus the analysis on the size range where periodic signals are most reliably detected, as longer eccDNAs are less frequent, likely reflecting reduced circularization efficiency, and exhibit increased variability that progressively obscures detectable periodic structure in the length distribution.

The spectral representation is dominated by low-frequency components associated with the global shape of the eccDNA length distribution. These components can obscure biologically relevant periodic signals at shorter length scales. To mitigate this effect, we first applied a high-pass cutoff of 0.005 using scipy.signal.butter to suppress very low-frequency variations in the global distribution, hereby removing periodicities longer than approximately 200 bp and enhancing the detection of biologically relevant signals at shorter length scales, including nucleosome organization signal.

To further isolate short-range periodic structure, we increased the high-pass cutoff to 0.05, removing periodicities longer than approximately 20 bp and thereby attenuating nucleosome-associated signals. This enhanced the relative contribution of higher-frequency components associated with DNA helical periodicity (10.5 bp).

The Butterworth filter was chosen for its maximally flat passband response, avoiding distortions in the retained frequency range. Its transfer function is defined as:

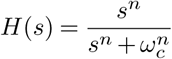

where *s* = *jω* is the complex frequency variable, *n* is the filter order controlling the slope, and *ω_c_*is the cutoff frequency. Frequencies below *ω_c_* are progressively attenuated, allowing us to focus on high-frequency signals corresponding to shorter and biologically meaningful frequencies.

To standardize the amplitude values obtained from the STFT and enable meaningful comparisons across different length segments and frequency components, we applied z-score normalization. This process involves subtracting the mean and dividing by the standard deviation of the amplitude values within each frequency band. Formally, for each frequency *f* , the normalized value *z* of an amplitude *A* is calculated as:

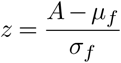

where *µ_f_* and *σ_f_* are the mean and standard deviation of amplitudes at frequency *f* across all time segments. This normalization removes scale differences and centers the data around zero with unit variance, highlighting relative deviations from the average signal strength. As a result, biologically relevant local high-frequency signals stand out more clearly against background fluctuations.

### Length distributions

In this study, we analysed eccDNA length distributions using two complementary smoothed representations: a global length distribution and a peak-centred local distribution. Raw eccDNA lengths were binned at single–base-pair resolution to generate discrete histograms representing length frequencies, which were subsequently smoothed using a one-dimensional Gaussian filter to reduce stochastic noise (**Figure 1D, E**).

The global distribution, evaluated over a fixed range up to 1000 bp, captures the overall eccDNA size landscape, which is dominated by nucleosome-associated organization (180 bp). To emphasize these large-scale structural features and enable robust identification of dominant modes, we applied a stronger Gaussian smoothing (σ = 3). We detected local maxima using scipy.signal.find_peaks, and the dominant peak was defined as the maximum of the smoothed global distribution.

To investigate fine-scale structure embedded within this nucleosome-dominated regime, we defined a 100-bp window centred on the dominant peak and reanalysed the distribution using weaker Gaussian smoothing (σ = 1). This local representation preserves short-range variability while remaining constrained within the nucleosome-associated length scale, allowing exploration of potential lower periodicity patterns consistent with DNA helical geometry periodicity (10.5 bp).

### Mapping nucleosomes

We obtained genome-wide nucleosome positioning data from Gaffney and colleagues (GSE36979; Downloaded from the Gene Expression Omnibus), comprising MNase-seq data from seven human lymphoblastoid cell lines (Gaffney et al., 2012). We converted The WIG track to BigWig format using UCSC tools v2.9 (Kent et al., 2010) and the chromosome sizes of the hg18 genome.

We called nucleosome dyads called from the MNase midpoint signal using a local-enrichment approach, similar to the analysis performed in the original research (Gaffney et al., 2012). For computational efficiency, each chromosome was processed in 5-Mb chunks. To ensure that scores near chunk edges were computed with complete windows, each chunk was extended by 1 kb on either side; dyads called within these flanking regions were discarded to avoid duplicate calls. This procedure yields the same results as processing each chromosome as a whole. We converted missing or NaN values within each chunk to 0. For every genomic position (i), We quantified MNase signal within two sliding windows: a proximal window of 15 bp (corresponding to a 31-bp window) and a broader background window of 100 bp (corresponding to a 201-bp window). We then calculated a score:

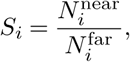

where 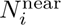 and 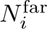 denote the number of MNase fragment midpoints within the near and far windows, respectively. Because the near window is contained within the far window, *S_i_* ranges from 0 to 1 and represents the fraction of midpoints in the 201-bp window that fall within the central 31 bp. Thus, *S_i_* measures the local enrichment of MNase fragment midpoints relative to the surrounding background, and we used it as a positioning score to identify candidate nucleosome dyad positions. We defined candidate nucleosomes as contiguous genomic regions with *S_i_ >* 0.4. Under a uniform distribution of midpoints, the expected value of *S_i_* is 31*/*201 *≈* 0.15; this threshold therefore requires an approximately 2.6-fold local enrichment over a non-positioned background. From each candidate region, we selected as the dyad the position with the highest positioning score; when several positions shared the maximum score, we selected the one closest to the center of the region. We discarded candidates with fewer than 10 midpoints in the far window. Calls separated by less than 147 bp were considered overlapping; in each overlapping group, we retained the call with the highest positioning score, breaking ties by the far-window count. We flagged calls as strong when *S_i_ ≥* 0.5 (approximately 3.2-fold enrichment). This procedure resulted in 4,100,086 dyad calls in the hg18 genome. We lifted over these calls to hg38 using the UCSC liftOver tool (v2.9) (Hinrichs et al., 2006), losing 321 calls. The final set contained 4,099,765 hg38 dyad calls, of which 3,158,492 were classified as strong. Only strong calls were used in downstream analyses.

### Nucleosome-associated positioning of eccDNA breakpoints

The breakpoint-positioning analysis was restricted to the four human datasets. Coordinates for *Hs-cancer_cells-MD* were called directly against GRCh38 by the nf-core/circdna pipeline, whereas the three Circle-Seq datasets (*Hs-blood-CS*, *Hs-muscle-CS*, *Hs-sperm-CS*) were obtained from the original publications in hg19 and lifted over to hg38 using the UCSC liftOver tool (Hinrichs et al., 2006), which failed to map 76 of 22,398 intervals (0.34%). Pooling the four datasets yielded 615,856 eccDNA intervals, which were not deduplicated across datasets.

We defined intergenic eccDNA by intersecting the pooled intervals against all gene-level features of the GENCODE v50 annotation (Frankish et al., 2023) using bedtools intersect (Quinlan and Hall, 2010), discarding any eccDNA overlapping an annotated gene by at least one base pair. Because the annotation includes all GENCODE gene biotypes, not only protein-coding genes, this is a conservative definition. Of the 615,856 pooled intervals, 134,551 were retained as intergenic.

To assess whether eccDNA breakpoints preferentially occurred at positions relative to nucleosome dyads, we identified the closest consensus nucleosome dyad to each circle breakpoint. We obtained the consensus dyads from the Gaffney et al. (2012) nucleosome positioning dataset described above and restricted them to strong calls (*S_i_ ≥* 0.5; 3,158,492 dyads). For each eccDNA, we considered both the start and end coordinates as breakpoints and independently identified the closest consensus dyad for each breakpoint. For each breakpoint, we calculated the signed distance between the nearest nucleosome dyad and the breakpoint (dyad position minus breakpoint position), such that positive values indicate dyads located downstream of the breakpoint.

To minimize potential confounding from differences coming from two contiguous nucleosomes, we restricted the breakpoint-positioning analysis to circles of up to 200 bp, leading to 34,435 out of the total 134,551 circles.

We calculated breakpoint-to-dyad distances for the observed eccDNA set and for 1,000 randomized eccDNA coordinate sets. Randomized coordinates were generated using bedtools shuffle, retaining each eccDNA on its original chromosome and excluding ENCODE blacklist regions (Amemiya et al., 2019). For each distance (*d*), the observed count of breakpoint–dyad pairs was divided by the corresponding count in each randomized dataset. These ratios were then averaged across the 1,000 randomizations to obtain the mean observed-to-expected enrichment profile that are described in all the subsequent analyses in this section.

We evaluated the resulting distance profile over distances of up to 500 bp from the nearest dyad. To visualize the overall nucleosome-associated pattern, we smoothed the profile using a centred 21-bp rolling mean. We calculated a 95% empirical confidence interval independently at each distance from the distribution of observed-to-expected ratios across permutation datasets and smoothed the confidence limits using the same procedure. We visualized the nucleosomal region as the canonical nucleosome-associated region extending approximately 83.5 bp on either side of the dyad, thereby accounting for a nucleosome length of 167 bp.

To examine fine-scale breakpoint positioning relative to the rotational organization of nucleosomal DNA, we restricted to positions within 60-bp of the dyad and smoothed it using a centred 5-bp rolling mean. We defined positions corresponding to successive approximately 10-bp intervals around the dyad to distinguish the two rotational orientations of the DNA minor groove. Specifically, we classified positions at approximately *±*5, *±*15, *±*25, *±*35, *±*45, *±*55, and *±*65 bp as minor-groove-inward-oriented positions, whereas we classified positions at approximately *±*0, *±*10, *±*20, *±*30, *±*40, *±*50, and *±*60 as minor-groove-outward-oriented positions (Noll, 1974; Luger et al., 1997; Richmond and Davey, 2003). For each class, we extracted enrichment values within a 5-bp window centred on the corresponding positions.

To isolate the fine-scale oscillatory component from the broader nucleosome-positioning profile, we detrended the observed-to-expected enrichment profile by subtracting a broad baseline estimated using a second-order Savitzky-Golay filter (Savitzky and Golay, 1964) with a 21-bp window from a locally smoothed profile generated using a 5-bp window. The 5-bp window was used to retain the fine-scale groove-in/groove-out oscillations, whereas the broader 21-bp window was used to estimate the underlying nucleosome-associated enrichment trend. Subtracting the broad profile from the locally smoothed profile therefore removed the gradual nucleosome-positioning effect while preserving the shorter 10-bp periodic component. We subsequently smoothed the resulting residual profile using a Gaussian filter (*σ* = 1.5 bp).

To compare minor-groove-inward and minor-groove-outward orientations, nucleotides were classified according to their proximity to the groove positions defined above. Nucleotides located within *±*2.5 bp of a minor-groove-inward position were classified as minor-groove-inward, whereas those within *±*2.5 bp of a minor-groove-outward position were classified as minor-groove-outward. We then compared residual enrichment values at minor-groove-inward- and outward-oriented positions using a two-sided Mann-Whitney U test. We assessed normality using the Shapiro-Wilk test and, when both distributions were consistent with normality, used Welch’s two-sample *t*-test instead. We reported effect sizes as Cohen’s *d* for the Welch test and rank-biserial correlation for the Mann-Whitney test. Because this comparison is made between positions of the smoothed enrichment profile rather than between independent circles, neighbouring positions are correlated by construction, and the resulting *n* (60 inward- and 55 outward-oriented positions) should be interpreted accordingly.

### Circle length phase relative to the 10.5-bp helical repeat

To assess whether eccDNA circle sizes showed a preferential phase relative to the canonical 10.5-bp helical repeat of B-DNA, we restricted the analysis to circles with a length of up to 200 bp. For each circle, we calculated its position within a 10.5-bp cycle from its length (*L*). We calculated the phase as the remainder of the circle length after division by 10.5 bp, normalized to the 10.5-bp period and expressed in degrees:

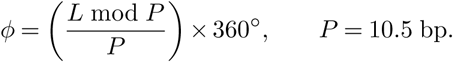

Because eccDNA lengths are integers and 10.5 bp corresponds to half-integer steps, phase values can take only 21 discrete values, spaced 360*^◦^/*21 *≈* 17.1*^◦^* apart; circles whose lengths differ by a multiple of 21 bp (two helical turns) are therefore assigned exactly the same phase. We visualized the number of circles at each of these 21 phase values across the 10.5-bp cycle as a histogram.

### Software, statistics, and visualization methods

All analyses were performed using Python 3.11. Signal-processing and statistical analyses were performed using SciPy (Virtanen et al., 2020). STFT analysis was performed using scipy.signal.stft, and Butterworth filtering was performed using scipy.signal.butter. Length-distribution peaks were identified using scipy.signal.find_peaks. One-dimensional Gaussian smoothing was performed using scipy.ndimage.gaussian_filter1d, and Savitzky-Golay filtering was performed using scipy.signal.savgol_filter (Savitzky and Golay, 1964). Rolling-window smoothing and tabulation of analysis results were performed using pandas. Statistical analyses were performed using scipy.stats, including the Shapiro-Wilk test, Welch’s two-sample *t*-test, and Mann-Whitney U test. Graph visualizations were generated using matplotlib (Hunter, 2007) and seaborn (Waskom, 2021). Software requirements and analysis code are available in the corresponding repository. Genomic interval operations and the generation of randomized coordinate sets were performed using bedtools v2.31.1 (Quinlan and Hall, 2010). BigWig signal was read using pyBigWig. Note that the random stream of bedtools shuffle is not guaranteed to be stable across bedtools versions, so permutation-derived values may differ marginally when reproduced with a different version.

## Data availability

All the data generated is available at Zenodo (Zabala et al., 2026a).

## Code availability

All the code is available at Zenodo (Zabala et al., 2026b) and https://github.com/ZabalaAitor/eccDNA_periodicity.

## Acknowledgments

AZ gratefully acknowledges the support and hospitality of the host institutions: Rigshospitalet Copenhagen University Hospital, Center for Genomic Medicine, and the University of Copenhagen, Center for Health Data Science. All circular detection analyses were conducted on the Hyperion cluster at the Donostia International Physics Center (DIPC). The authors acknowledges the technical and human support provided by the DIPC Supercomputing Center. The authors have used generative AI technology to improve the readability and overall quality of the manuscript. The use of AI has not altered the content or messages conveyed in the manuscript, focusing solely on refining the writing for clarity and enhanced readability.

## Supplementary Information

**Figure S1.**
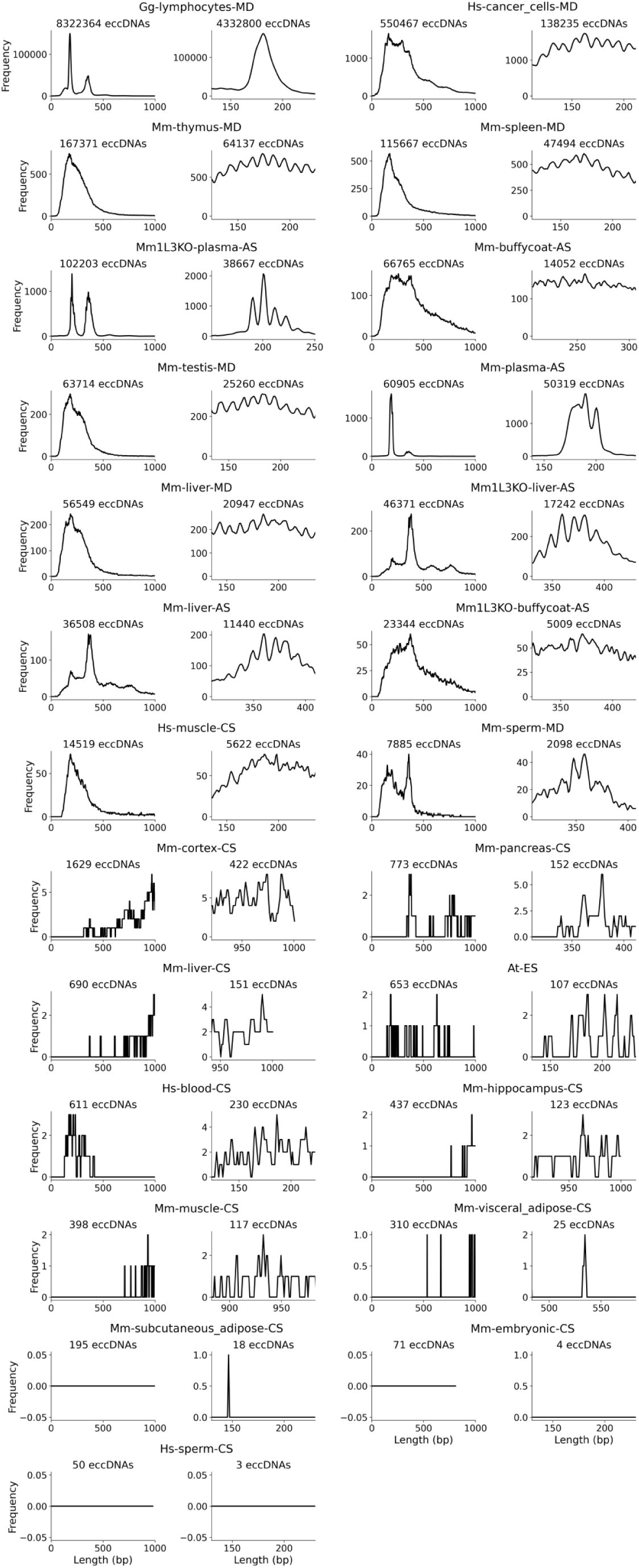
eccDNA length distribution. Left panels display the global length distributions of eccDNA length distributions up to 1000 bp, showing dominant and secondary peaks across dataset. Right panels zoom into a narrower window around the dominant peak to highlight subtle short-length oscillatory patterns.

**Figure S2.**
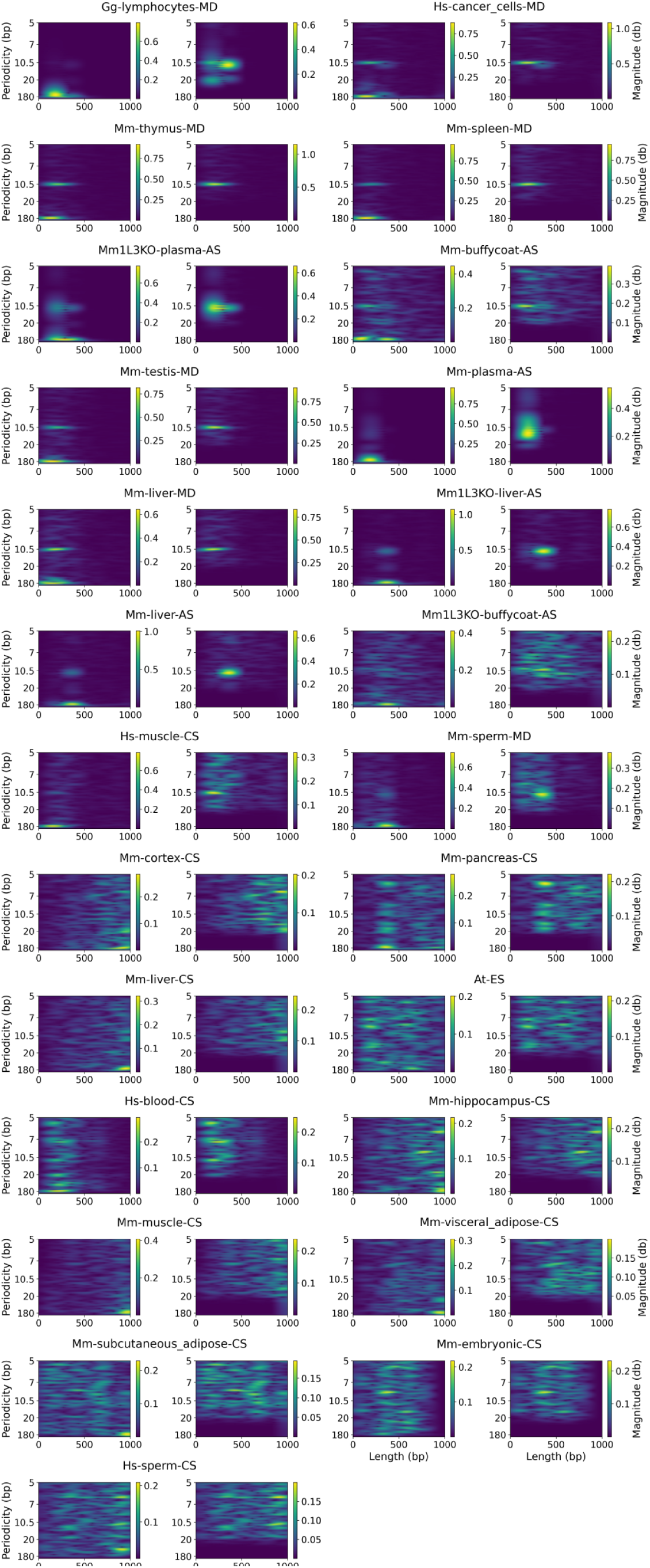
Short-Time Fourier Transform (STFT) analysis. Left panels display global periodicity profiles obtained using a Butterworth high-pass filter with a cutoff frequency of 0.005, capturing the 180 bp periodicity (low-frequency signal) Right panels focus exclusively on the local periodicity using a higher cutoff frequency of 0.05, capturing the 10.5 bp periodicity (high-frequency signal).

**Figure S3.**
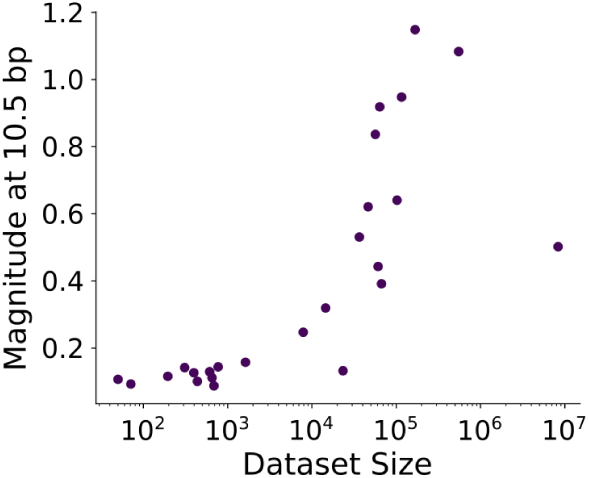
Effect of dataset size on detection of 10.5 bp periodicity in eccDNA. Relationship between the magnitude of the 10.5 bp periodic signal (high-frequency signal) and the number of eccDNA molecules detected per dataset. Each point represents a dataset; the x-axis (log scale) shows sample size and the y-axis shows spectral magnitude at 10.5 bp.

**Figure S4.**
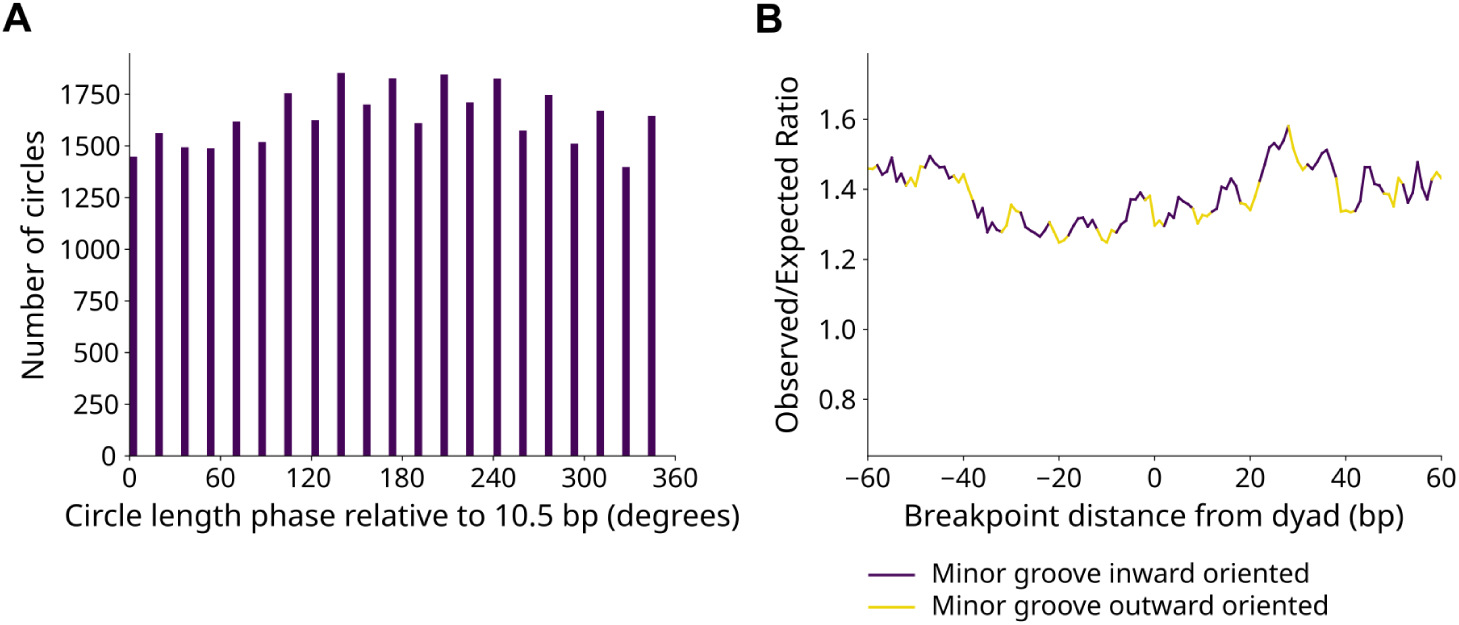
Zoomed-in breakpoint enrichment around consensus nucleosome dyads. Zoomed view of the observed/expected breakpoint enrichment profile around consensus nucleosome dyads, highlighting the fine-scale periodic pattern within *±*60 bp of the dyad. Breakpoint enrichment shows recurrent peaks at positions corresponding to outward- and inward-oriented minor grooves, revealing the periodic structure that is less apparent in the full nucleosomal profile.

